# The *Fusarium graminearum* transporters Abc1 and Abc6 are important for xenobiotic resistance, trichothecene accumulation, and virulence to wheat

**DOI:** 10.1101/2021.06.15.448535

**Authors:** Sean P. O’Mara, Karen Broz, Yanhong Dong, H. Corby Kistler

## Abstract

The plant pathogenic fungus *Fusarium graminearum* is the causal agent of Fusarium Head Blight (FHB) disease on small grain cereals. *F. graminearum* produces trichothecene mycotoxins such as deoxynivalenol (DON) that are required for full virulence. DON must be exported outside the cell to cause FHB disease, a process that may require the involvement of membrane-bound transporters. In this study we how the deletion of membrane-bound transporters results in reduced DON accumulation as well as reduced FHB symptoms on wheat. Deletion of the ATP-Binding Cassette (ABC) transporter *Abc1* results in the most severe reduction in DON accumulation and virulence. Deletion of another ABC transporter, Abc6, also reduces FHB symptoms to a lesser degree. Combining deletions fails to reduce DON accumulation or virulence in an additive fashion, even when including an *Δabc1* deletion. Heterologous expression of *F. graminearum* transporters in a DON-sensitive strain of yeast confirms Abc1 as a major DON resistance mechanism. Yeast expression further indicates that multiple transporters, including Abc1 play an important role in resistance to the wheat phytoalexin 2-benzoxazolinone (BOA) and other xenobiotics. Thus, Abc1 may contribute to wheat virulence both by allowing export of DON and by providing resistance to the wheat phytoalexin BOA. This research provides useful information which may aid in designing novel management techniques of FHB or other destructive plant diseases.

## Introduction

Fusarium is a cosmopolitan genus comprised of not only soil-borne and endophytic fungi which live asymptomatically within hosts, but also fungal species that are economically and agronomically important plant pathogens (Imazaki and Kadota 2015; Lofgren et al. 2018; Wachowska et al. 2017; Waweru et al. 2014). *Fusarium graminearum* is one of the causal agents of Fusarium Head Blight (FHB) disease of small grain cereals and has received considerable attention due to its ability to act as a plant pathogen and to produce mycotoxins that impact animal and human health. While occurring world-wide, FHB has been a persistent problem in the United States for many years, especially since a large outbreak in the 1990’s. The disease management costs and direct impact of FHB outbreaks have resulted in annual losses exceeding $1.4B in the United States (Wilson et al. 2018).

*F. graminearum* produces several secondary metabolites which act as virulence factors during plant host infection (Bahadoor et al. 2018; Proctor 1995; Wipfle et al. 2019), including sesquiterpenoid trichothecenes. Trichothecenes inhibit protein synthesis by binding to the peptidyl-transferase domain of eukaryotic ribosomes (Fried and Warner 1981; Garreau de Loubresse et al. 2014; Harris and Gleddie 2001), an effect which poses significant risk to plants and animals. The major trichothecene produced by many strains of *F. graminearum* is deoxynivalenol (DON), which is essential for full virulence on wheat (Desjardins et al. 1996) and may persist as a contaminant in affected grains. Exposure to DON can lead to many cytological effects in eukaryotes, including DNA/RNA synthesis disruption, ribotoxic stress, induction of apoptosis, and membrane cytotoxicity (Pestka 2007; Rocha et al. 2005). Export of trichothecences by *F. graminearum*, whether by vesicular traffic or by membrane-bound transporters, may be essential for full virulence to plants (Abou Ammar et al. 2013; Gardiner et al. 2013; Menke et al. 2012; O’Mara et al. 2020) Additionally, due to potential trichothecene toxicity, *F. graminearum* must either sequester or export this secondary metabolite or risk self-inhibition (Menke et al. 2012; Wang et al. 2018).

One of the primary sources of resistance to toxic secondary metabolites in fungi is through the actions of membrane-bound transporters, such as ATP-binding cassette (ABC) and Major-facilitator superfamily (MFS) transporters (Gulshan and Moye-Rowley 2007; Perlin et al. 2014). The typical structure of an ABC transporter includes two core domains, a nucleotide-binding domain (NBD) and a transmembrane domain consisting of six transmembrane-spanning helices (TMD) (Kovalchuk and Driessen 2010). MFS transporters are generally smaller than ABC transporters and consist of 12 or 14 transmembrane spanning helices without any NBD domains. ABC transporters utilize adenosine triphosphate (ATP) to move molecules across membranes while MFS transporters utilize electrochemical membrane potential (often due to a pH gradient) to accomplish the same functionality (Coleman and Mylonakis 2009).

Both ABC and MFS transporters have many crucial functions in fungi including nutrient uptake/acquisition, waste removal, cellular signaling, self-defense, and other developmental processes. In pathogenic fungi, transporters are important for fungal infection, colonization, and disease progression (Perlin et al. 2014). The first identified pleiotropic drug resistance protein, the ABC-G transporter *PDR5*, was described from baker’s yeast *Saccharomyces cerevisiae* (Gulshan and Moye-Rowley 2007). *PDR5* has orthologs in most other ascomycetes including Fusarium where it has been associated with virulence by conferring resistance to plant phytoalexins (Coleman et al. 2011; Fleissner et al. 2002).

*F. graminearum* has 62 predicted ABC proteins (Kovalchuk and Driessen 2010), four of which appear to be involved in virulence (Yin et al. 2018). The ABC transporter FgAbc9 (FGSG_07325) plays a crucial role in the export of the plant defense hormone salicylic acid, and is also involved in growth, resistance to the anti-fungal compound tebuconazole, and accumulation of DON (Qi et al. 2018). Perhaps the most studied membrane-bound transporter of *F. graminearum*, the ABC transporter Abc1 (FGSG_04580) also is important for resistance to fungicides and other xenobiotics. Deletion mutants of *Abc1* show reduced virulence during wheat crown and root rot infections (Gardiner et al. 2013). Similarly, other *F. graminearum* strains with the FGSG_04580 gene deleted (called *Abc3* in this reference), show reduced virulence on wheat, barley, and maize, as well as altered mycotoxin production (Abou Ammar et al. 2013). Abc1 is also involved in zearalenone (an estrogenic metabolite of *F. graminearum*) transport and tolerance to antifungal compounds (Abou Ammar et al. 2013; Gardiner et al. 2013; Lee et al. 2011).

In addition to ABC transporters, a number of *F. graminearum* MFS transporters have been suggested to be involved in toxin export and resistance. Wang and colleagues (2018) identified 33 transporters, 15 of which were MFS transporters, which were specifically induced by externally added DON. Furthermore, a number of the identified MFS transporters had homologs known to be involved in resistance to toxic compounds and secondary metabolite export in other fungi (Wang et al. 2018). Within the core trichothecene biosynthetic gene cluster of *F. graminearum* is the gene *Tri12* (FGSG_03541) which encodes an MFS transporter (Proctor et al. 2009). Disruption of the *Tri12* gene of *F. sporotrichioides*, a close relative of *F. graminearum*, results in highly reduced trichothecene accumulation in culture and reduced resistance to exogenously applied trichothecenes (Alexander et al. 1999). However, disruption of *Tri12* in *F. graminearum* results in only slightly reduced DON accumulation (Menke et al. 2012).

The goal of this research was to test other ABC and MFS transporters or combinations of transporters, for their role in trichothecene export and xenobiotic resistance in *F. graminearum*. In addition to further characterization of the transporters Tri12 and Abc1, we included other ABC and MFS transporters previously identified and suspected to have a role in DON export. The ABC transporter *Abc6* (FGSG_11028) and an MFS transporter *Mfs1* (FGSG_07802) are either regulated by or otherwise co-expressed with genes under the control of the trichothecene regulatory protein Tri6 (Seong et al. 2009). During *F. graminearum* infection of wheat, *Abc6* and *Mfs1* are significantly up-regulated contemporaneously with high levels of DON production (Zhang et al. 2012) and in a *Tri12* deletion mutant background, these two genes are significantly upregulated under toxin producing conditions (Nakajima et al. 2015). We hypothesize that their gene products may play a redundant role in DON transport, and “fill in” when Tri12 is not available.

We hypothesize that multiple, functionally redundant mechanisms for DON transport may exist within *F. graminearum* cells. If this is the case, deletion of only a single mechanism for transport may not lead to a discernable phenotype with respect to fungal growth, DON accumulation, or pathogenic aggressiveness. However, disruption of two or more mechanisms for DON transport might be expected to have greater impact. To test this, we combined deletion mutations for different membrane bound transporters to assess the effect of disrupting multiple export pathways. These deletion mutants were analyzed for growth and DON accumulation *in planta*. Additionally, the protein coding regions of individual genes were expressed in a DON sensitive strain of *S. cerevisiae* to verify their direct involvement in DON export and resistance to inhibition by a number of xenobiotic compounds.

## Materials and Methods

### Generating F. graminearum deletion mutants

Genetic deletion mutants of *F. graminearum* strain PH-1 (NRRL 31084) were generated using the split marker homologous recombination technique (Goswami 2012) with modifications previously described (O’Mara et al. 2020). A previously generated *Δtri12* disruption mutant was used (Menke et al. 2012). Primers utilized in the transformation and confirmation process are listed in Table S1. Target genes were replaced with the neomycin phosphotransferase (NPT) or nourseothricin acyltransferase (NAT) resistance genes as selectable markers (Fuchs et al. 2004; Menke et al. 2013). Transformations and single-spore isolations were completed as previously described (O’Mara et al. 2020). Up to 10 transformant colonies were picked for further single-spore isolation and genetic confirmation. Mutants of *F. graminearum* were crossed to generate double- and triple knockout mutants. Mutant parents contained different antibiotic-resistance selectable markers. Sexual crosses and ascospore collection and screening were performed as previously described (Pasquali and Kistler 2006; O’Mara et al. 2020). Single spore isolation of each transformant was performed as for single knockout mutants, except that media contained two antibiotics.

### Confirmation of F. graminearum transformants

Site-directed deletion of native genes in *F. graminearum* mutants was confirmed by PCR as previously described (O’Mara et al. 2020) using the CTAB genomic DNA extraction technique (Gale et al. 2011). Site-directed gene deletion and replacement with an antibiotic-resistance selectable marker was confirmed by amplifying the targeted locus using a primer pair which annealed upstream and downstream of the gene. Changes in amplicon size corresponding to the replacement of the native gene with the antibiotic-resistance marker indicated site-directed gene replacement. In cases where the amplicon sizes were too similar to distinguish using this method, an either/or amplification technique was employed. In the latter case, a forward primer which annealed upstream of the targeted locus was paired with either a reverse primer which annealed to the end of the native gene or a reverse primer which annealed to the end of the antibiotic-resistance gene. This amplification pair would indicate whether the full native gene remained in its resident locus, or if the full antibiotic-resistance gene was inserted into the locus.

### Growth of F. graminearum in culture

Conidial suspensions of each *F. graminearum* mutant were made by growing in 50 mL of carboxy-methyl cellulose (CMC) medium (Cappellini and Peterson 1965) for ∼5 days. Conidia were collected and enumerated as previously described (O’Mara et al. 2020) then suspended in water at 2 × 10^4^ conidia/mL for *in vitro* analyses or 1 × 10^6^ conidia/mL for plant inoculations.

To assess the ability of *F. graminearum* deletion mutants to utilize different laboratory media, deletion mutants were grown on carrot, ½ PDA, Czapek-Dox, complete, minimal, and V8 agar media (Klittich and Leslie 1988; Puhalla and Spieth 1983; Rodriguez Estrada et al. 2011). A 3 mm diameter plug of mycelium was placed at the center of each and grown in triplicate at 25°C with 12 h light, 12 h dark for 3 days before determining the area (mm^2^) of hyphal growth using a Carestream 4000MM Pro Image Station, with Carestream Molecular Imaging Software v.5.2.2.15761 (Carestream Health, Inc., Rochester, NY, USA).

### DON accumulation and pathogenicity of F. graminearum transporter deletion mutants

The ability of *F. graminearum* mutants to cause FHB symptoms and accumulate DON was evaluated in wheat cultivar Norm, as previously described (O’Mara et al. 2020). Two weeks after point-inoculation of the fifth fully formed spikelet, heads were scored for FHB disease symptoms (bleached, shriveled, necrotic grains) by counting the number of diseased spikelets, up to 10, surrounding the point of inoculation. The inoculated spikelet was then removed from the wheat head and weighed in a tared 1-dram screw-cap vial, frozen at -80°C overnight and analyzed for DON, 3-ADON, and 15-ADON (total DON) by GC-MS using methods previously described (Goswami and Kistler 2005).

### Generation of F. graminearum transporter containing yeast transformants

*F. graminearum* transporters were expressed in the DON-sensitive *Saccharomyces cerevisiae* strain YZGA515 (Poppenberger et al. 2003), generously provided by Dr. Gerhard Adam. Expression plasmids were generated using Gateway cloning technology (Invitrogen, Thermo-Fisher Scientific, Waltham, MA, USA). A cDNA of the *F. graminearum Abc1* gene codon-optimized for yeast was synthesized by Invitrogen GeneArt gene synthesis and inserted onto the pENTR221 vector. Codon-optimized cDNAs for *Tri12, Mfs1*, and *Abc6* were synthesized by Integrated DNA Technologies (IDT, Coralville, IA, USA) and inserted on pUC-IDT vectors. *E. coli* strains containing the empty expression vector ZM552 or the yeast pleiotropic drug resistance transporter Pdr5 on the ZM552 plasmid were purchased from the DNASU plasmid repository (Arizona State University, Tempe, AZ, USA). ZM552 was used as an empty vector control for all tests, and Pdr5 was used as a positive control complementation.

To begin the Gateway cloning process, forward primers containing the attB1 site and reverse primers containing the attB2 site (Table S1) were purchased from Invitrogen. These primers were used to amplify the *F. graminearum* transporter coding regions (*Tri12, Mfs1*, and *Abc6*) from the pUC-IDT plasmids. Once purified, the attB site flanked amplicons were cloned onto the Gateway donor vector pDONR221 using the Gateway BP Clonase II enzyme mix kit, generating pENTR221 vectors. Expression plasmids were generated by cloning the *F. graminearum* transporter coding regions from the pENTR221 vectors (now *Tri12, Abc1, Mfs1*, and *Abc6*) onto the ZM552 empty vector using the Gateway LR Clonase II enzyme mix kit. Following the Gateway LR cloning step, all the transporter genes from *F. graminearum* and *S. cerevisiae* were located on the ZM552 vector backbone and named pEXP552_*Gene Name* (e.g. pEXP552_FgTri12 or pEXP552_ScPdr5). All pENTR221 and pEXP552 plasmids were cloned into Invitrogen OmniMAX 2-T1 or New England Biolabs (Ipswich, MA, USA) 10-Beta *E. coli* for propagation. The pDONR221 plasmid was cloned into Invitrogen ccdB Survival 2-T1 *E. coli* for propagation. Confirmations of all cloning steps were performed by extracting plasmid DNA from *E. coli* using the Qiagen Miniprep kit (Hilden, Germany) and performing double restriction enzyme digests using EcoRI and HindIII purchased from New England Biosciences (NEB, Ipswich, MA, USA), using the manufacturer’s recommended procedures.

All transformation plasmids were transformed into *S. cerevisiae* YZGA515 using the Sigma-Aldrich (St. Louis, MO, USA) YEAST1 yeast transformation kit, using manufacturer’s recommended procedures. Transformed YZGA515 strains were streaked and maintained on Synthetic Complete Medium (Dunham et al. 2015) supplemented with leucine drop out powder (Sigma Aldrich). To confirm proper transformation of yeast strains, plasmid DNA was extracted from transformed yeast using the Qiagen Miniprep kit and re-transformed into *E. coli* for propagation. Plasmid DNA from re-transformed *E. coli* was extracted again using the Qiagen Miniprep kit and digested using EcoRI and HindIII as before. Restriction digests from original transformed *E. coli* strains were compared to digests from re-transformed *E. coli* to confirm identical plasmid composition.

### Expression of F. graminearum transporters in a susceptible yeast line

Transformed *S. cerevisiae* YZGA515 strains were analyzed for their sensitivity to DON and its acetylated derivatives 3-acetyl deoxynivalenol (3-ADON) and 15-acetyl deoxynivalenol (15-ADON). Pre-cultures of YZGA515 transformants were grown in liquid SCM-leu medium for 4-5 days at 30°C and 300 rpm. Pre-cultures were diluted with 2x SCM-leu to an optical density (600 nm absorbance) of 0.1 for inoculation. In a 96-well microtiter plate, 100 µL of YZGA515 inoculum was added to 100 µL ddH_2_O supplemented with a concentration gradient of DON, 3-ADON, or 15-ADON. Concentration series included 0, 20, 30, 60, 120, and 250 ppm DON or 3-ADON, and 0, 5, 10, 15, 20, and 30 ppm 15-ADON. All six YZGA515 transformants were inoculated onto the same 96-well microtiter plate containing six DON/ADON concentrations, with an extra well containing 1x SCM-leu without yeast as a medium control. Plates were sealed with parafilm and incubated in the dark for 5 days at 30°C and 300 rpm. After incubation, well contents were homogenized by gentle pipetting and the optical density of each well was taken using a Thermo-Fisher Scientific Varioskan Flash with SkanIt RE software. DON/ADON plates were run in triplicate and sensitivity was analyzed as a percent change in optical density compared to 0 ppm control.

To test whether low pH would better facilitate DON export in *S. cerevisiae* YZGA515 lines expressing *F. graminearum* MFS transporters (Gardiner et al. 2009), the plate assay was repeated using glycine-HCl buffered SCM-leu. To make buffered medium, 1.875 g glycine was added to 200 mL 2x SCM-leu. Mixture was then titrated with 5M HCl to pH 2.5. The buffered medium was then brought up to 250 mL using ddH_2_O and sterilized by filtering through a Corning 0.22 µm filter unit (Corning Life Sciences, Corning, NY, USA). Assay setup and optical density measurements were performed as previously described above.

To understand how the *F. graminearum* transporters may provide resistance to xenobiotic compounds, the transformed yeast lines were tested against a concentration series of the plant phytoalexin 2-benzoxazolinone (BOA). Transformant strains were grown in 5 mL SCM-leu medium for 4∼5 days at 30°C and 300 rpm. Cultures were centrifuged at 1,200 x g for 5 minutes and resuspended in sterile water to an optical density of OD=0.38. BOA was dissolved in DMSO to concentration of 500 mg/mL. In a 96-well plate, 4 µL of YZGA515 transformant cell suspensions were inoculated into 200 µL SCM-leu medium containing 0, 100, 200, 300, 400, or 500 ppm BOA, with a final concentration of 0.1% DMSO. After inoculation, 96-well plates were incubated at 30°C and 300 rpm for 3 days. Optical density of each well was determined using a Thermo-Fisher Scientific Varioskan Flash with SkanIt RE software. Plates were run in triplicate and sensitivity was analyzed as a percent change in optical density compared to 0 ppm control.

The yeast transformants were also tested against 24 xenobiotic compounds at 4 concentrations using Biolog plate PM24C (Biolog Inc, Hayward, CA, USA). Transformants were grown and resuspended as for BOA plates then 630 µL of the cell suspension was added to 29.38 mL of Biolog PM inoculating fluids using the manufacturer’s recommended protocol and mixed thoroughly. Using an 8-channel pipette, 100 µL of the inoculation fluid was transferred into each well of the 96-well PM24C plate. Plates were incubated at 30°C for 3 days before recording the OD value of each well. Plates were conducted in triplicate for each yeast transformant. Raw OD values for each transformant were subtracted from the OD values of the ZM552 empty vector control for each replicate. Afterwards, the relative OD values for each transformant were averaged across the three replicates for a final growth value.

### Data analysis

*F. graminearum* growth rates, FHB disease symptoms, and DON concentrations were analyzed in R Statistical Software version 3.5.1 (R Core Team 2018). Data were analyzed by a one-way ANOVA with a Tukey’s post-hoc test to compare all pair-wise interactions. ANOVAs for *in vitro* and *in planta* inoculations were blocked for inoculation date, to better address inter-genotype differences. Sensitivity of S. *cerevisiae* to exogenous DON and BOA was analyzed by a one-way ANOVA with a Dunnett’s post-hoc test (Hothorn et al. 2008) to compare differences from the control group. ANOVAs were conducted on raw optical density values (600nm) while figures show percent growth of genotype compared to the 0 ppm concentrations.

## Results

### Confirmation of F. graminearum mutants

Genes for *F. graminearum* membrane-bound transporters potentially involved in DON export and FHB virulence were knocked out to assess their function in these processes. All knockouts were performed by replacing the native gene with a neomycin (neoR) or nourseothricin (ntcR) resistance gene. Mutant genotypes were confirmed by PCR, using changes in amplicon size or by selectively amplifying either the wild-type locus or its resistance-construct replacement (Figure S1). All transformations yielded at least one mutant which grew on antibiotic selection and amplified a PCR product indicative of a successful site-directed, gene replacement deletion; most yielded multiple deletion candidates and a single mutant was selected for further analysis. Deletion phenotypes were further confirmed for DON accumulation by comparing neoR or ntcR deletion mutants with an independently generated hygromycin (hygR) deletion mutant. All deletion mutant pairs (neoR vs. hygR or ntcR vs. hygR) accumulated the same levels of DON *in vitro* (Figure S2).

### F. graminearum transporter deletion mutants are not altered in growth or morphology

Deletion mutants of *F. graminearum* were assessed for their growth rates and phenotype on six different laboratory media. Growth of all mutant genotypes at three days post inoculation was not significantly different from the wildtype (p>0.05; Table S2). Additionally, deletion of membrane-bound transporters in *F. graminearum* did not manifest any overt phenotypes when grown on laboratory media, even when combining multiple deletions (Figure S3).

### Multiple transporter mutants show reduced DON accumulation and virulence in wheat

*F. graminearum* knockout mutants were tested for their ability to accumulate DON plus its acetylated derivatives (total DON) and cause FHB symptoms *in planta*. Numerous combinations of transporter deletion mutants were significantly reduced in the ability to accumulate DON (F=35.76, df=346, p<2.2e-16). Two levels of effect were seen (Figure 1A). Deletion of the *Tri12* gene reduced DON accumulation in wheat approximately 35% while deletion of the *Abc1* transporter gene reduced DON accumulation in wheat by 65%. Combining the *Δabc1* and the *Δtri12* alleles did not reduce DON accumulation beyond that seen for the *Δabc1* alone. Likewise, combining the *Δabc6* or *Δmfs1* alleles with *Δtri12*, as double or triple mutants, didn’t reduce DON accumulation beyond the reduction seen in the *Δtri12* mutant alone.

**Figure 1:**
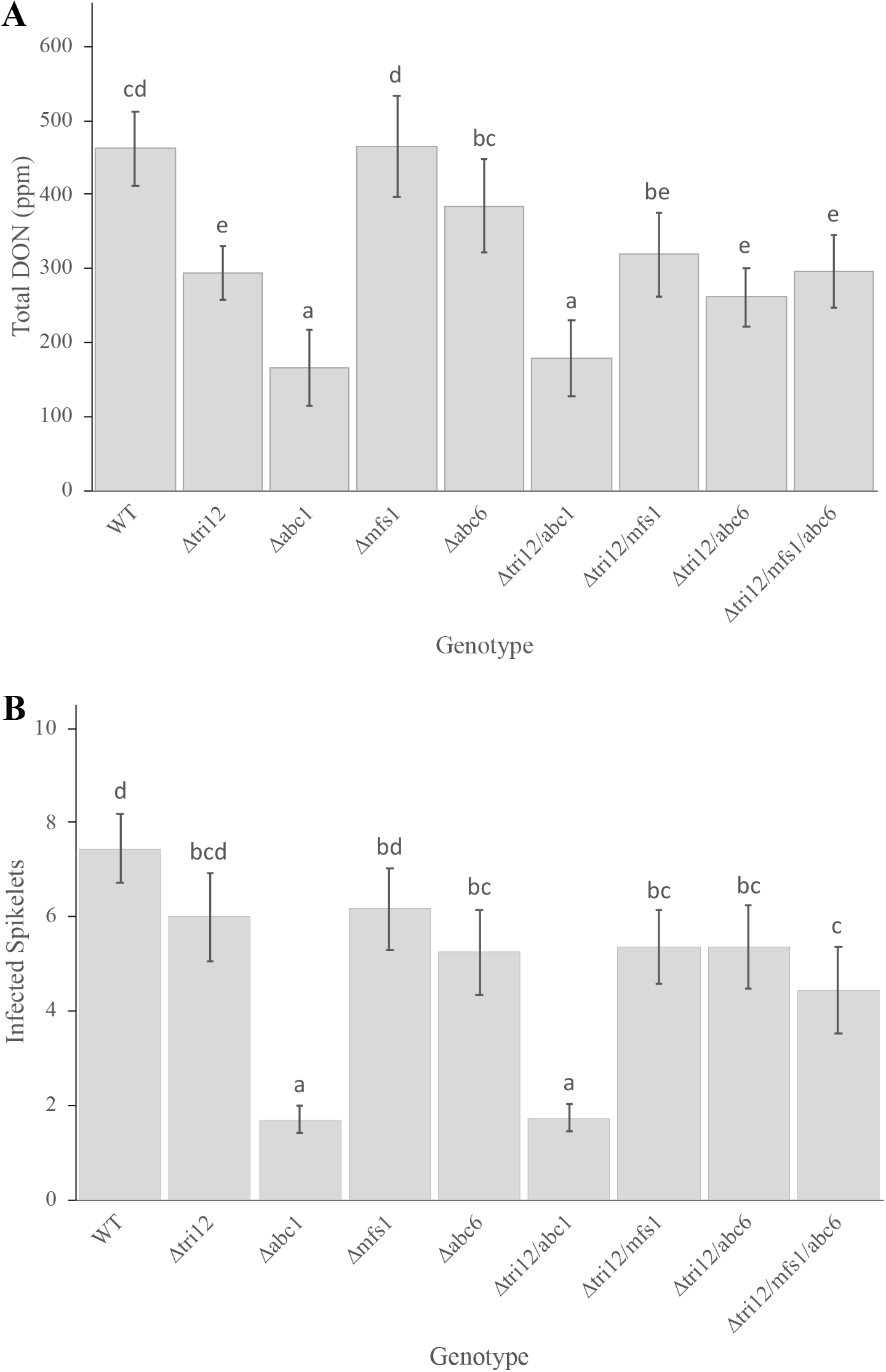
Total DON accumulation *in planta* and pathogenicity of *F. graminearum* transporter mutants. Total DON accumulation (A) and disease progression (B) 14 days post inoculation (n=40). Mean + 95% CI. Data analyzed by one-way ANOVA. Genotypes with the same letter are not significantly different as determined by a Tukey’s post hoc analysis.

Infected spikes were assessed 14 days post inoculation for the spread of FHB disease symptoms and significant differences were detected between the genotypes (F=25.96, df=346, p<2.2e^-16^). FHB symptoms followed a somewhat similar trend as seen for DON accumulation *in* planta (Figure 1B). However, deletion of the Abc6 gene reduced FHB spread by ∼30% while deletion of A*bc1* reduced FHB spread by ∼78%. Again, combining multiple mutations was not able to significantly reduce FHB symptoms below those of the parent genotypes. These results fail to indicate an additive effect of multiple transporter deletions but do suggest that Abc1 plays a substantial role in DON accumulation and FHB virulence, while other transporters play a more minor role.

### F. graminearum transporters alter DON and BOA sensitivity in a susceptible yeast line

Four *F. graminearum* membrane-bound transporters were expressed in a DON-sensitive yeast strain YZGA515 and exposed to a concentration series of DON and acetylated derivatives 15-ADON and 3-ADON to assess the function of these transporters. The yeast multidrug resistance transporter *Pdr5* expressed in YZGA515 was used as a positive control, and YZGA515 without an insert was used as a negative control. The YZGA515 *FgAbc1* strain showed a substantial increase in resistance to DON, 15-ADON, and 3-ADON (Figure 2), showing up to six-fold increased resistance to the toxins as compared to the vector control. The native *ScPdr5* strain also showed a substantial increase in resistance to the toxins. The others, including the *FgTri12, FgMfs1*, and *FgAbc6* strains, showed little to no increase in resistance to DON and 15-ADON but *FgAbc6* may allow for some resistance to higher levels of 3ADON. Interestingly, the two MFS transporter strains, *FgTri12* and *FgMfs1*, were actually more sensitive to DON and 15-ADON at lower concentrations than the empty vector control. To determine if these MFS transporters, which are predicted proton antiporters, would provide resistance to DON at low pH, all transformants were subsequently tested against DON in acidified medium.

**Figure 2:**
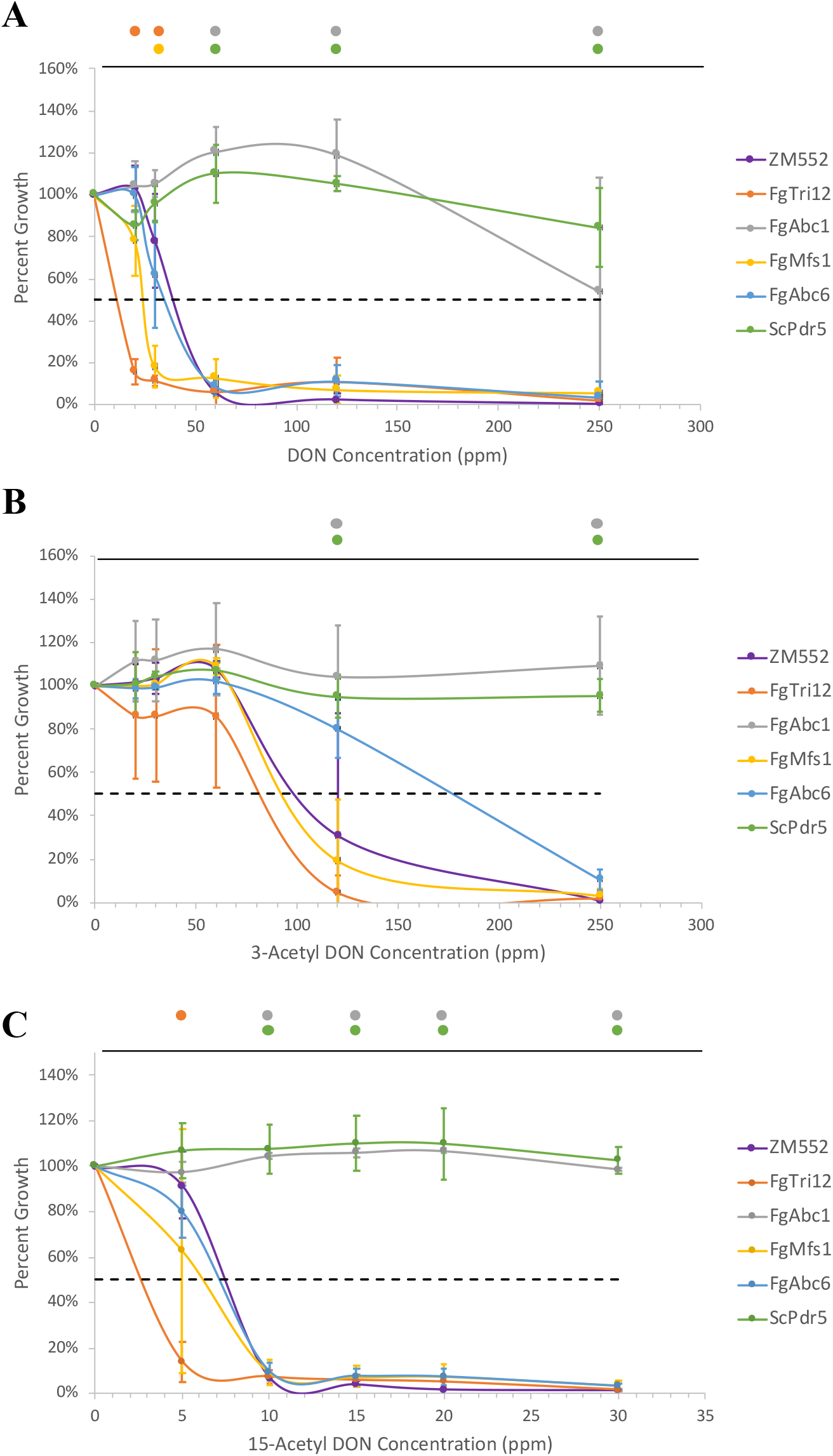
Growth of yeast YZGA515 transformants exposed to exogenous DON and ADON. Growth of *S. cerevisiae* strain YZGA515 expressing *F. graminearum* transporters after 5 days exposure to exogenous (A) DON, (B) 3-ADON, and (C) 15-ADON. Measurements are the average (n=3) percent growth compared to the 0 ppm concentration ± 95% confidence interval. Dashed black line indicates EC50. Colored dots above graph indicate genotypes which are significantly different (p<0.05) from the ZM552 control at each concentration. One-way ANOVA conducted on raw optical density (600 nm) measurements at each concentration.

When the assays were repeated in medium buffered to pH 2.5 with glycine-HCl, the *FgTri12* strain was increased in resistance and was no more sensitive to DON than the empty vector control at 20 ppm, the lowest concentration of DON tested (Figure 3). However, at the next higher concentration (30 ppm) the *FgTri12* strain again showed greater DON sensitivity than the control as did *FgMfs1* and *FgAbc6*. This suggests that the hydrogen ion concentration can impact the activity of *FgTri12* but only at lower DON concentrations. This is of note because *F. graminearum* establishes an acidic extracellular environment under toxin-inducing conditions (Gardiner et al. 2009) that may provide the proton-motive force to affect Tri12-mediated transport. The *FgAbc1* and *ScPdr5* strains continued to show increased resistance to DON as compared to the empty vector control (Figure 3).

**Figure 3:**
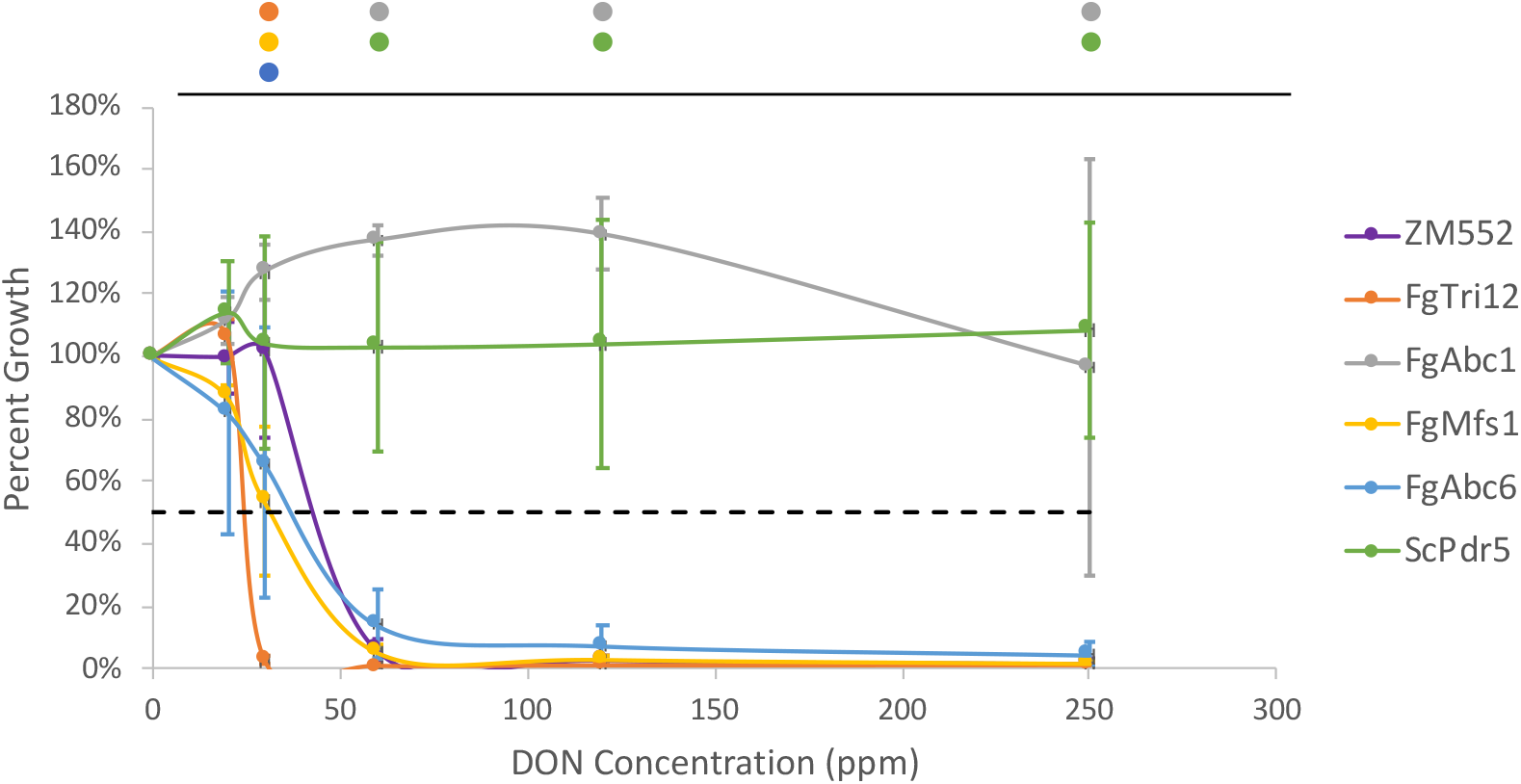
Growth of yeast YZGA515 transformants exposed to exogenous DON at pH 2.5. Growth of *S. cerevisiae* strain YZGA515 expressing *F. graminearum* transporters after five days exposure to exogenous DON/ADON at pH 2.5. Measurements are the average (n=3) percent growth compared to the 0 ppm concentration ± 95% confidence interval. Dashed black line indicates EC50. Colored dots above graph indicate genotypes which are significantly different (p<0.05) from the ZM552 control at each concentration. One-way ANOVA conducted on raw optical density (600 nm) measurements at each concentration.

To assess the ability of the *F. graminearum* transporters to provide resistance to xenobiotics, YZGA515 strains were exposed to a concentration series of the wheat phytoalexin 2-benzoxazolinone (BOA) for 3 days. The empty vector control strain ZM552 showed moderate sensitivity to BOA, with an estimated EC50 value of approximately 250 ppm (Figure 4). The *FgMfs1* strain showed increased sensitivity to BOA, with an estimated EC50 value of 160 ppm. Surprisingly, the remaining strains (*FgTri12, FgAbc1, FgAbc6*, and *ScPdr5*) all showed increased resistance to BOA, with all having an estimated EC50 value of 350 ppm.

**Figure 4:**
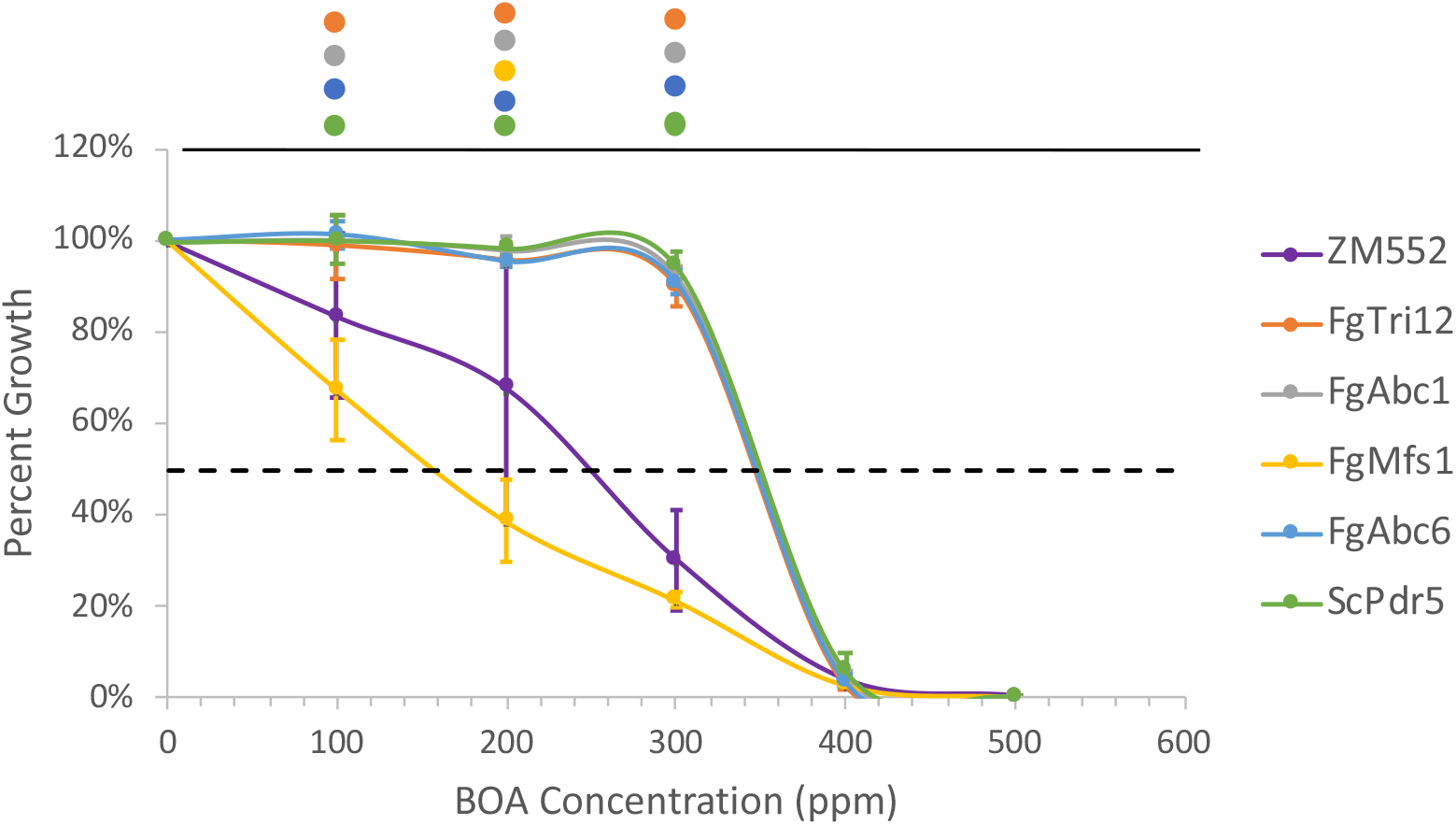
Growth of yeast YZGA515 transformants exposed to exogenous BOA. Growth of *S. cerevisiae* strain YZGA515 expressing *F. graminearum* transporters after three days exposure to the plant phytoalexin 2-benzoxazolinone (BOA). Measurements are the average (n=3) percent growth compared to the 0 ppm concentration ± 95% confidence interval. All *F. graminearum* transporter transformants showed increased resistance to BOA, except for the FgMfs1 lines which was more sensitive. Dotted black line indicates EC50. Colored dots above graph indicate genotypes which are significantly different (p<0.05) from the ZM552 control at each concentration. One-way ANOVA conducted on raw optical density (600 nm) measurements at each concentration.

Chemical sensitivity was further analyzed by exposing the yeast strains to 24 xenobiotics included in the Biolog PM24C plate (Biolog Inc, Hayward, CA, USA) (Figure S4). With the exception of a marked increased resistance to azole fungicides in the *ScPdr5* positive control and the *FgAbc1* strain, there was only moderate resistance to xenobiotics for the other transformants. Other than *FgMfs1*, all strains showed moderately increased resistance to antibiotics and metal salts with the *FgAbc1* and *ScPdr5* strains additionally having increased resistance to anticancer and other antifungal compounds. This finding fits into a larger pattern in which the *F. graminearum* Abc1 transporter plays a key role in the export of endogenous trichothecenes and a broad range of exogenous xenobiotics, including azole fungicides (Abou Ammar et al. 2013; Gardiner et al. 2013; Lee et al. 2011), providing both increased virulence and defense for *F. graminearum*.

## Discussion

While ABC- and MFS transporters play multiple roles in the normal physiology of fungi, for pathogenic species, they may also allow resistance to stresses related to host infection and can be essential for full expression of animal and plant pathogenesis (Cavalheiro et al. 2018; Coleman and Mylonakis 2009). Additionally, transporters are known to confer resistance to man-made antifungal compounds with consequences in clinical and agricultural settings (Ma and Michailides 2005; Sanglard 2016).

Transporters also can be important for the export of fungal secondary metabolites. These metabolites are typically produced by gene clusters encoding enzymes for the biosynthesis of the metabolites themselves as well as transporters presumed to allow for metabolite secretion. When these metabolites are mycotoxins, export mediated by gene cluster-encoded transporters has been suggested to be important to avoid self-inhibition. However, deletion mutants for the transporters in mycotoxin gene clusters sometimes can secrete nearly wild-type levels of the toxin and show little or no adverse impact on growth during toxin-inducing conditions (Chang et al. 2004; Proctor et al. 2003). This has led to the idea that additional secretion mechanisms may be operative within the cell that also function to export toxins (O’Mara et al. 2020). For the trichothecene biosynthetic gene cluster in *F. graminearum*, deletion of the cluster-associated MFS transporter, Tri12, indeed has been shown to have minimal effect on secretion of DON and related metabolites (Menke et al. 2012). Deletion analysis of an ABC transporter Abc1 and components of vesicular transport pathways each suggested that alternate pathways for DON secretion were possible (O’Mara et al. 2020). Here we sought to test the impact of mutation on multiple transporters in *F. graminearum*, alone and in combination, on pathogenicity to wheat and DON accumulation *in planta*.

None of the transporter deletion mutants used in this study presented altered growth or macroscopic changes in phenotype when tested on laboratory media (Table S2 and Figure S3) indicating that the individual and combined effect of these transporters on the normal vegetative growth of *F. graminearum* is minimal. This suggests that these transporters may be associated with other functions such as the transport of secondary metabolites, which by definition are only produced during limited parts of the life cycle and are not necessary for primary metabolic function of an organism (Keller et al. 2005). While Tri12 is encoded within the trichothecene biosynthetic gene cluster, genes for the *Mfs1* and *Abc6* lie within the biosynthetic genes clusters for other known natural products; the mycotoxin Fusarin C and the siderophore Malonichrome, respectively (Sieber et al. 2014; Oide et al. 2014). *Abc1* is located near (∼25 kb) a predicted polyketide synthase, *Pks29*, and an O-methyltransferase (Sieber et al. 2014), and has been suggested to be involved in the transport of numerous *F. graminearum* secondary metabolites including trichothecenes and zearalenone (Abou Ammar et al. 2013; Lee et al. 2011). Taken together, there is correlative evidence that these transporters may be involved to varying degrees in secondary metabolite transport but not in primary vegetative growth.

By expressing these transporters in yeast, we sought to directly test their ability to transport DON and related compounds by measuring the ability of strains to overcome DON toxicity. Yeast strains expressing Fusarium transporters were also tested for sensitivity to a panel of chemicals including the wheat phytoalexin BOA, which may impact pathogenicity toward wheat (Kettle et al. 2015). The yeast strain YZGA515 is deficient in all major multidrug resistance transporters (Poppenberger et al. 2003) and is completely inhibited by 10 ppm 15-ADON and 60 ppm DON (Figure 2), the major trichothecenes produced by the wild type strain used in this study. As predicted, the *F. graminearum* Abc1 transporter performed nearly as well as its yeast ortholog, the Pdr5 multidrug resistance transporter, for increasing resistance of YZGA515 to DON, 15-ADON, and 3-ADON (Figure 2). The other *F. graminearum* transporters tested were not able to increase resistance to DON or 15-ADON although Abc6 may confer resistance at higher levels of 3-ADON. Surprisingly, the MFS transporters Mfs1 and especially Tri12, seemed to allow for greater sensitivity at DON and 15-ADON at lower concentrations.

Additionally, yeast strains expressing *FgTri12, FgAbc1, FgAbc6*, or *ScPdr5* provided similar levels of increased resistance to BOA (Figure 4). This observation is consistent with the idea that multiple *F. graminearum* MFS and ABC transporters are capable of transporting BOA and thereby conferring redundant modes of resistance to the phytoalexin. Previously, a *Δabc1* mutant of *F. graminearum* failed to show increased sensitivity to BOA *in vitro* (Gardiner et al. 2013). However, the concentration of BOA used in that study was equal to the highest concentration used in the present study (500 ppm), at which all strains tested here were sensitive. Additionally, several single ABC transporter mutants of *F. graminearum* did not grow at significantly different rates compared to wildtype on multiple concentrations of BOA (Abou Ammar et al. 2013). Based on our results in yeast, previously published results may have not detected an effect of individual transporters *in situ* due to functional redundancies of other MFS and ABC transporters.

Except for Abc1, there is a poor correspondence between the ability of the transporters to confer DON resistance in yeast and the ability contribute to DON accumulation by *Fusarium*. This may reflect a fundamental difference in the manner in which these transporters act in *Fusarium*. The other membrane-bound transporters studied here (Tri12, Mfs1, and Abc6) may be indirectly associated with DON accumulation, rather than more directly involved in DON export as with Abc1. While the ABC multidrug resistance transporters are expected to be targeted to the plasma membrane in yeast and *Fusarium* (Egner et al. 1995; Lee et al. 2011) we have previously noted that Tri12 in *Fusarium* localizes to motile vesicles that may fuse with the vacuole or the plasma membrane (Menke et al. 2012). Because Tri12 and Mfs1 expressed in yeast actually increases sensitivity to DON, our hypothesis is that Tri12 may facilitate DON uptake by these vesicles that may then be transported to the vacuole for sequestration or to the plasma membrane for export. Previously we noted that the plasma membrane localized SNARE protein Sso2, essential for subapical vesicular exocytosis, was required for wildtype DON accumulation (O’Mara et al. 2020). Moreover, combining mutations for *Abc1* and *Sso2* reduced DON accumulation in an additive manner indicating that both vesicular transport and direct export via the multidrug transporter Abc1 contributed to DON export in a non-redundant manner.

In conclusion the ABC transporter Abc1 plays a significant role in the export of the trichothecene DON both *in vitro* and *in planta* and may act as the primary membrane-bound transporter involved in DON export. Two other membrane-bound transporters studied here, Tri12 and Abc6, show significant involvement in DON accumulation either *in vitro* or *in planta* but may act indirectly in conjunction with vesicular transport mechanisms or by allowing for greater virulence necessary for maximum DON accumulation in wheat by transport of BOA or other small molecules. Disruption of *Abc1* through novel management techniques such as host-induced or spray-induced gene silencing (HIGS and SIGS respectively) (Koch et al. 2016; Qi et al. 2019), combined with other management techniques in an integrated pest management system, may provide better control of FHB.

## Acknowledgements

The authors greatly thank Dr. Gerhard Adam of the University of Natural Resources and Life Sciences, Vienna for generously providing *S. cerevisiae* strain YZGA515. The authors would also like to thank Dr. Fumiaki Katagiri of the University of Minnesota and Zixuan Zhong of the Research Center of Bioenergy and Bioremediation at Southwest University, Chongqing, P.R. China for providing helpful feedback in the development of this research. Funding was provided by United States Department of Agriculture, United States Wheat and Barley Scab Initiative, award FY18-KI-021 and award 2018-67013-28512 from the Agriculture and Food Research Initiative of the National Institute of Food and Agriculture, United States Department of Agriculture. Mention of trade names or commercial products in this article solely for the purpose of providing specific information and does not imply recommendation or endorsement by the United States Department of Agriculture (USDA). USDA is an equal opportunity provider and employer.

## Supplemental Information

**Figure S1:**
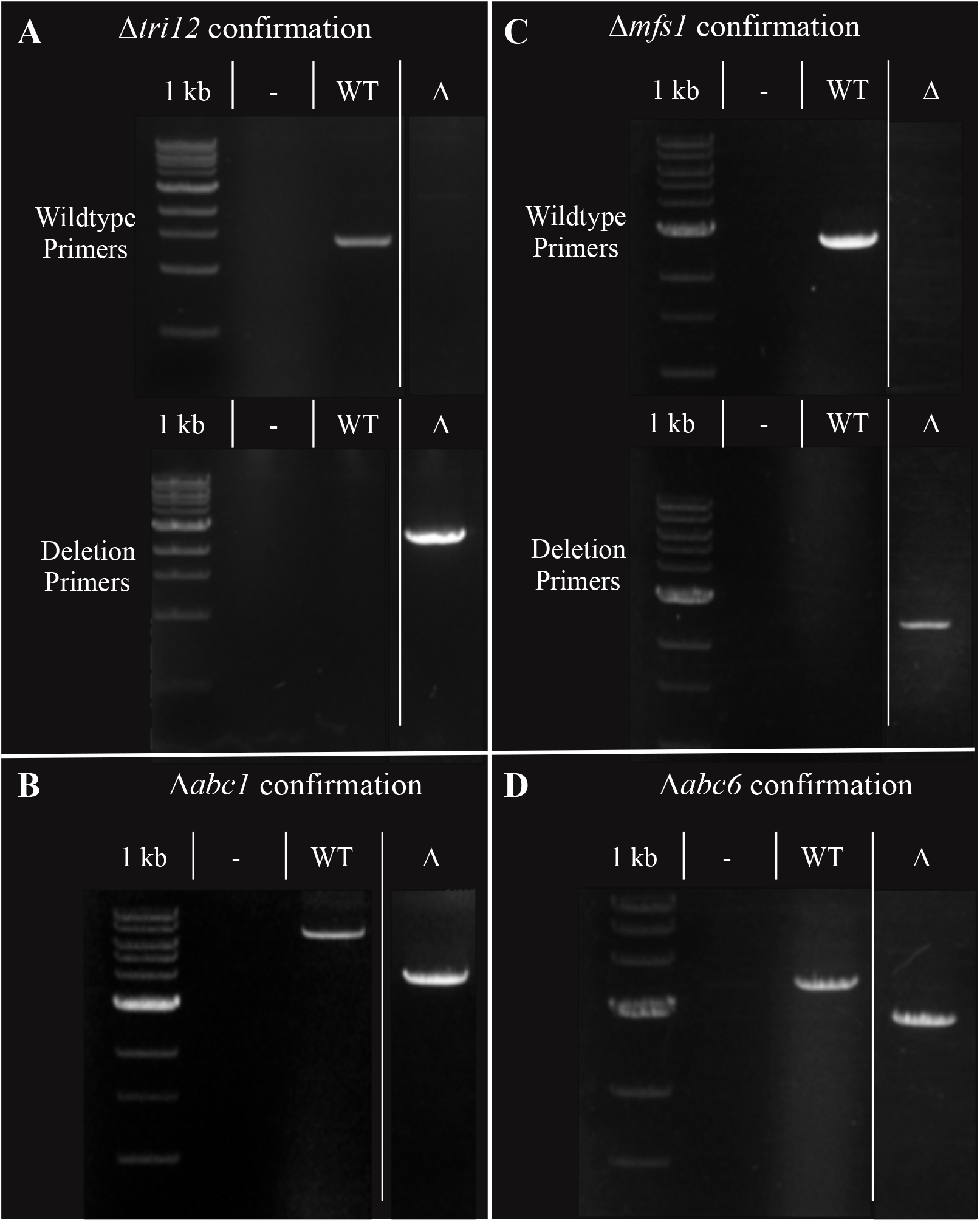
Confirmation PCR of *F. graminearum* deletion mutants. Size discrepancy or presence/absence determination of proper genetic deletions for (A) *Δtri12* knockout, (B) *Δabc1* knockout, (C) *Δmfs1* knockout, and (D) *Δabc6* knockout. Size discrepancy primers flank 5’ and 3’ of manipulated locus. Presence/absence wild type amplification primers bind upstream to manipulated locus and end of native gene; deletion amplification primers bind upstream to the target locus and end of antibiotic resistance gene. Vertical lines between lanes indicate removed lanes. 1 kb = 1 kb ladder, - = No DNA, WT = *F. graminearum* PH1 DNA, Δ = deletion mutant DNA.

**Figure S2:**
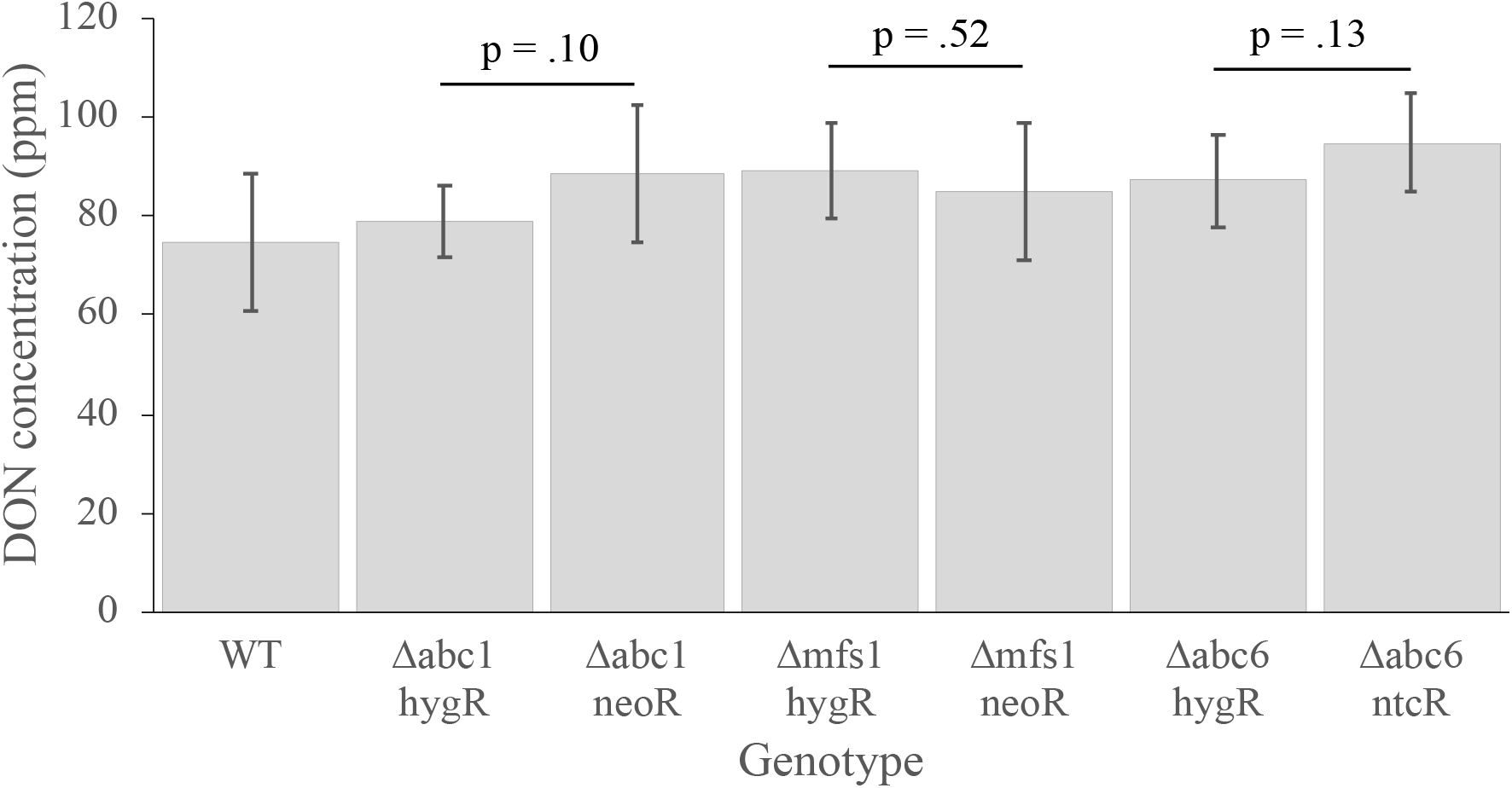
*In vitro* DON accumulation comparison of independently deleted *F. graminearum* transporters. Transporter deletions were conducted twice, using different antibiotic selectable markers, to confirm mutant phenotypes. Data analyzed by Student’s T-test. All comparisons between independent transformation mutants showed insignificant differences in DON accumulation.

**Figure S3:**
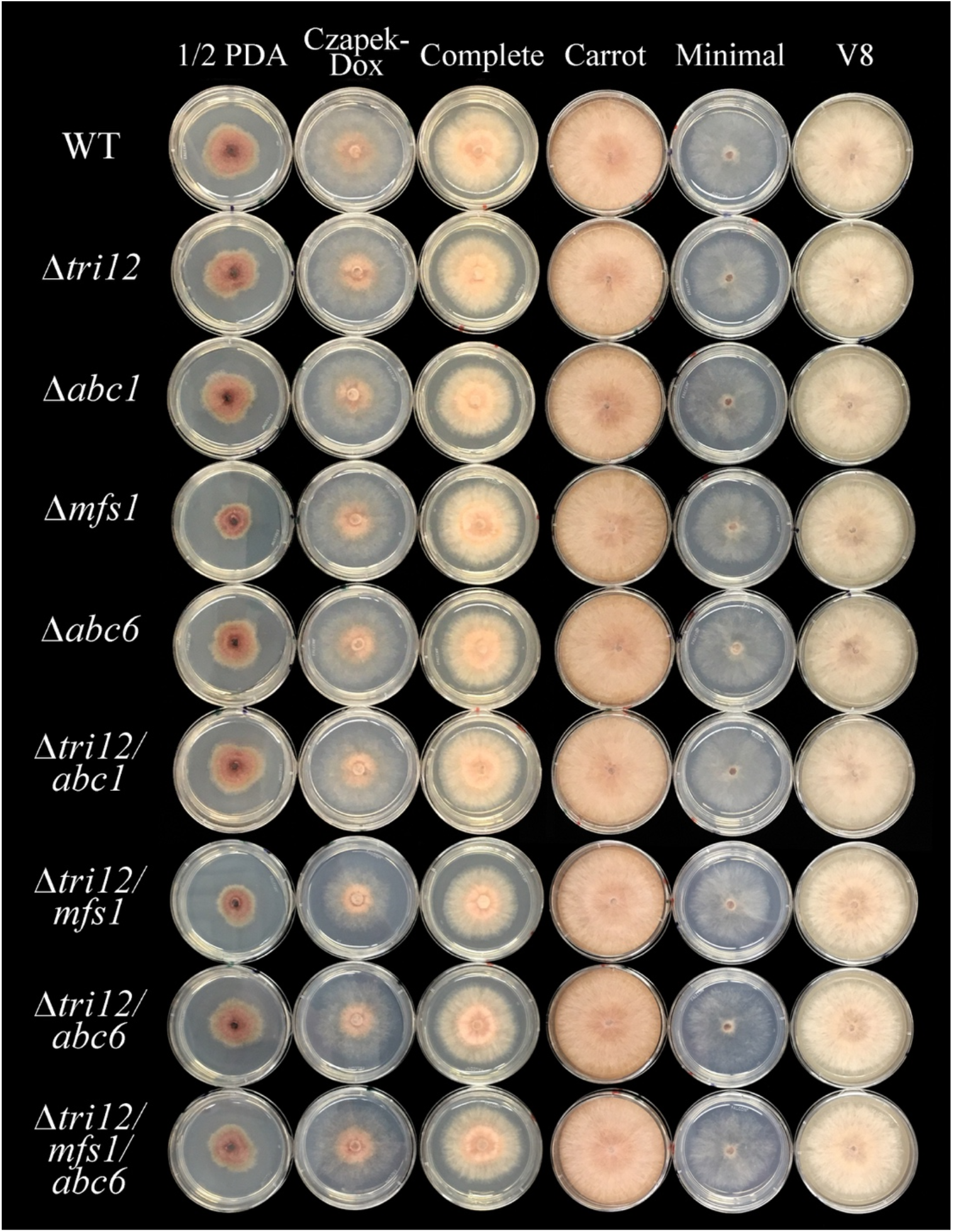
Phenotype of *F. graminearum* genotypes on laboratory media. Gene deletion did not result in any overt phenotypic changes in *F. graminearum* mutants, including multi-knockout mutants.

**Figure S4:**
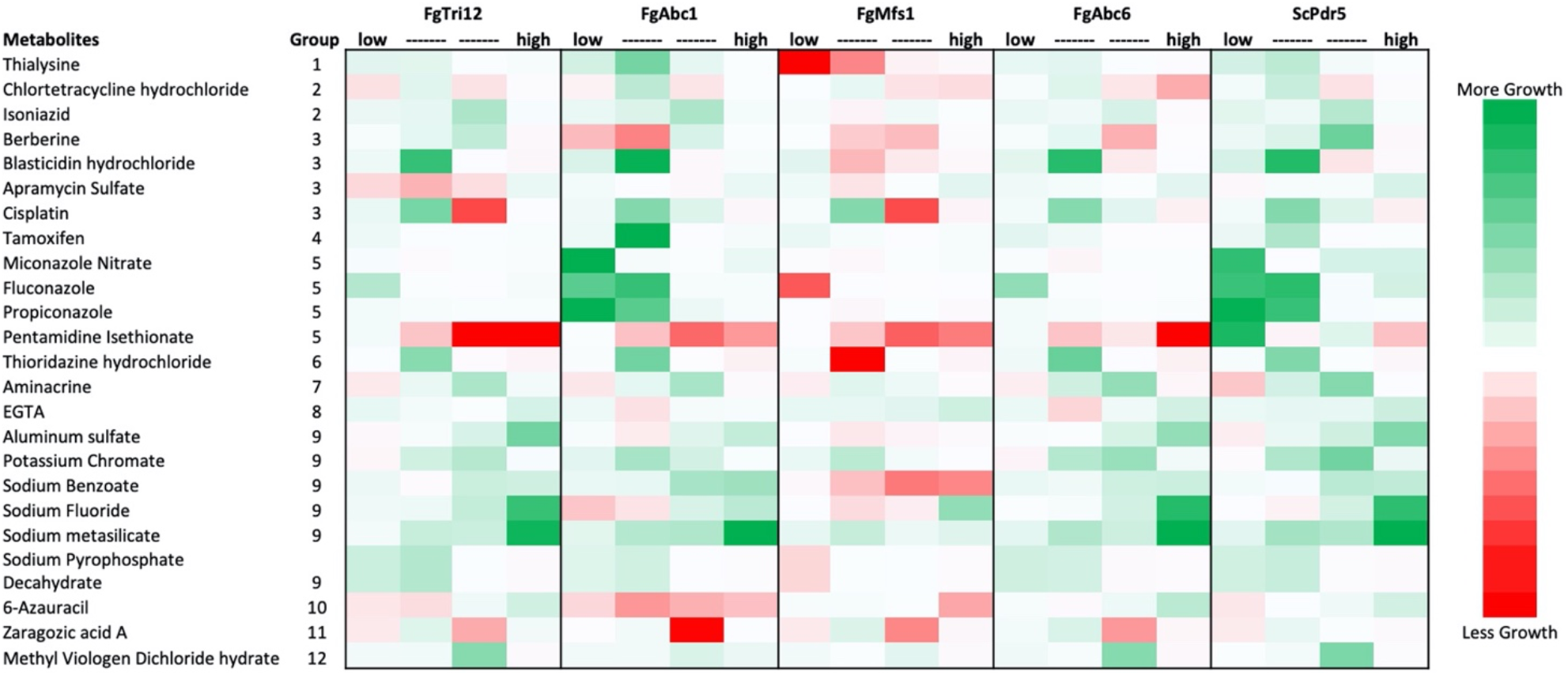
Growth of *F. graminearum* transporter expressing yeast on xenobiotics. Relative growth of transformed yeast YZGA515 expressing *F. graminearum* transporters compared to an empty vector control. Green cells indicated more growth of transformant compared to control; red cells indicate less growth. 1=Amino acid analog, 2=Antibacterial, 3=Antibiotic, 4= Anticancer, 5=Antifungal, 6=Antipsychotic, 7=Antiseptic, 8=Chelating Agent, 9=Metal salt, 10=Nucleotide analog, 11= Polyketide, 12=Viologen.

**Table S1:**
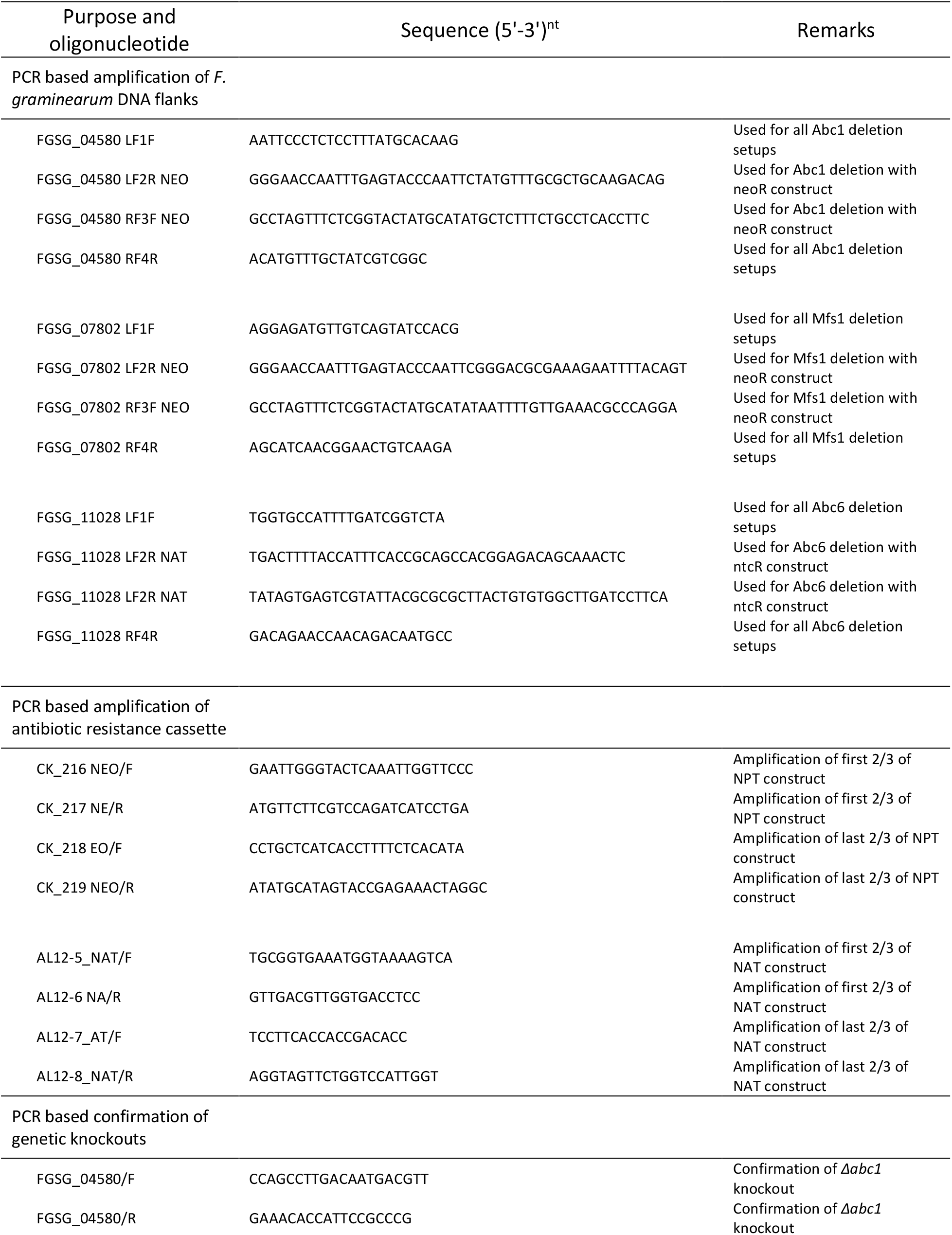
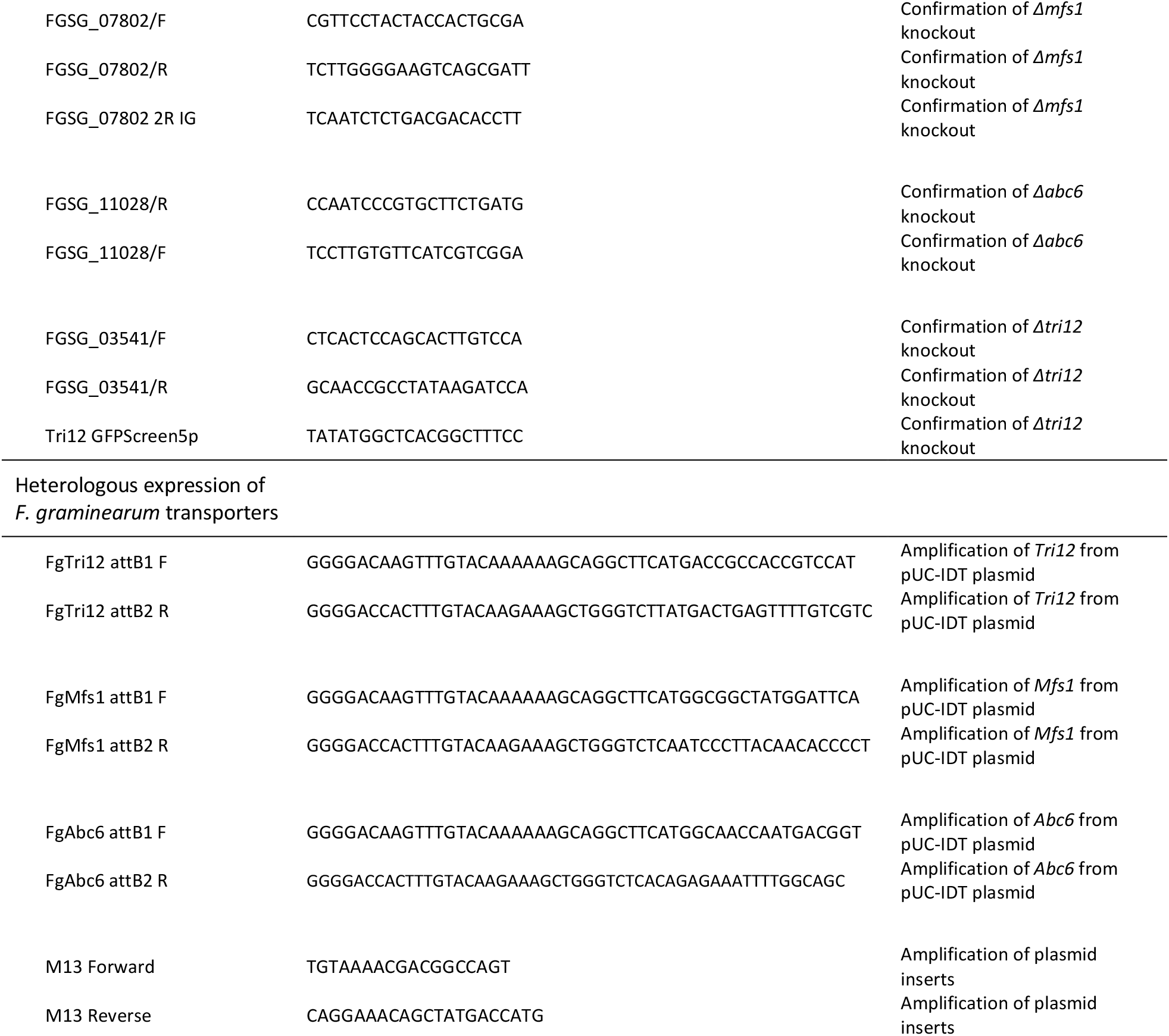
Oligonucleotide primers used for amplification, deletion, and confirmation of F. graminearum mutants.

**Table S2:** Colony area (mm^2^) of *F. graminearum* genotypes grown on laboratory media. Mean ± standard deviation (n=3) of *F. graminearum* colonies grown for 3 days on laboratory media. No significant differences in colony area were found between genotypes for each medium (p>0.05).

| Strain | Medium |  |  |  |  |  |
| --- | --- | --- | --- | --- | --- | --- |
|  | 1/2 PDA | Czapek-Dox | Complete | Carrot | Minimal | V8 |
| WT | 213 ± 52 | 886 ± 56 | 709 ± 104 | 2013 ± 100 | 830 ± 20 | 609 ± 49 |
| $\Delta tri12$ | 445 ± 204 | 1033 ± 94 | 843 ± 36 | 2031 ± 87 | 909 ± 75 | 585 ± 16 |
| $\Delta abc1$ | 358 ± 22 | 912 ± 83 | 753 ± 126 | 1929 ± 67 | 822 ± 76 | 530 ± 15 |
| $\Delta mfs1$ | 333 ± 26 | 1013 ± 118 | 812 ± 40 | 2027 ± 56 | 923 ± 51 | 555 ± 7 |
| $\Delta abc6$ | 327 ± 11 | 928 ± 128 | 794 ± 18 | 2062 ± 123 | 845 ± 57 | 551 ± 30 |
| $\Delta tri12/abc1$ | 200 ± 16 | 883 ± 81 | 710 ± 14 | 1923 ± 17 | 747 ± 30 | 579 ± 23 |
| $\Delta tri12/mfs1$ | 187 ± 44 | 823 ± 248 | 678 ± 228 | 1969 ± 29 | 798 ± 87 | 588 ± 71 |
| $\Delta tri12/abc6$ | 245 ± 105 | 944 ± 316 | 489 ± 187 | 1880 ± 58 | 807 ± 92 | 450 ± 133 |
| $\Delta tri12/mfs1/abc6$ | 266 ± 147 | 898 ± 282 | 714 ± 251 | 1942 ± 70 | 753 ± 54 | 561 ± 48 |

## Literature Cited

1. Abou Ammar, G., Tryono, R., Döll, K., Karlovsky, P., Deising, H. B., and Wirsel, S. G. R. 2013. Identification of ABC transporter genes of Fusarium graminearum with roles in azole tolerance and/or virulence. PLoS One. 8:e79042 Available at: http://www.pubmedcentral.nih.gov/articlerender.fcgi?artid=3823976&tool=pmcentrez&rendertype=abstract.

2. Alexander, N. J., McCormick, S. P., and Hohn, T. M. 1999. TRI12, a trichothecene efflux pump from Fusarium sporotrichioides: Gene isolation and expression in yeast. Mol. Gen. Genet. 261:977–984.

3. Bahadoor, A., Brauer, E. K., Bosnich, W., Schneiderman, D., Johnston, A., Aubin, Y., et al. 2018. Gramillin A and B: Cyclic lipopeptides identified as the nonribosomal biosynthetic products of Fusarium graminearum. J. Am. Chem. Soc. 140:16783–16791 Available at: http://pubs.acs.org/doi/10.1021/jacs.8b10017 [Accessed August 1, 2019].

4. Cappellini, R. A., and Peterson, J. L. 1965. Macroconidium formation in submerged cultures by a non-sporulating strain of Gibberella zeae. Mycologia. 57:962 Available at: http://www.jstor.org/stable/3756895?origin=crossref [Accessed March 26, 2018].

5. Cavalheiro, M., Pais, P., Galocha, M., and Teixeira, M. C. 2018. Host-pathogen interactions mediated by MDR transporters in fungi: As pleiotropic as it gets! Genes (Basel). 9 Available at: https://pubmed.ncbi.nlm.nih.gov/30004464/ [Accessed September 23, 2020].

6. Chang, P. K., Yu, J., and Yu, J. H. 2004. aflT, a MFS transporter-encoding gene located in the aflatoxin gene cluster, does not have a significant role in aflatoxin secretion. Fungal Genet. Biol. 41:911–920.

7. Coleman, J. J., and Mylonakis, E. 2009. Efflux in fungi: La piece de resistance. PLoS Pathog. 5.

8. Coleman, J. J., White, G. J., Rodriguez-Carres, M., and Vanetten, H. D. 2011. An ABC transporter and a cytochrome P450 of Nectria haematococca MPVI are virulence factors on pea and are the major tolerance mechanisms to the phytoalexin pisatin. Mol. Plant. Microbe. Interact. 24:368–376.

9. Desjardins, A. E., Proctor, R. H., Bai, G., McCormick, S. P., Shaner, G., Buechley, G., et al. 1996. Reduced virulence of trichothecene-nonproducing mutants of Gibberella zeae in wheat field tests. Mol. Plant-Microbe Interact. 9:775–781 Available at: http://www.apsnet.org/publications/mpmi/backissues/Documents/1996Abstracts/Microbe09-775.htm.

10. Dunham, M. J., Gartenberg, M. R., and Brown, G. W. 2015. Methods in yeast genetics and genomics : a Cold Spring Harbor Laboratory course manual. 2015 Editi. Cold Spring Harbor Laboratory Press.

11. Egner, R., Mahé, Y., Pandjaitan, R., and Kuchler, K. 1995. Endocytosis and vacuolar degradation of the plasma membrane-localized Pdr5 ATP-binding cassette multidrug transporter in Saccharomyces cerevisiae. Mol. Cell. Biol. 15:5879–5887.

12. Fleissner, A., Sopalla, C., and Weltring, K.-M. 2002. An ATP-binding cassette multidrug-resistance transporter is necessary for tolerance of Gibberella pulicaris to phytoalexins and virulence on potato tubers. Mol. Plant. Microbe. Interact. 15:102–108.

13. Fried, H. M., and Warner, J. R. 1981. Cloning of yeast gene for trichodermin resistance and ribosomal protein L3. Proc. Natl. Acad. Sci. U. S. A. 78:238–42 Available at: http://www.pubmedcentral.nih.gov/articlerender.fcgi?artid=319027&tool=pmcentrez&rendertype=abstract.

14. Fuchs, U., Czymmek, K. J., and Sweigard, J. A. 2004. Five hydrophobin genes in Fusarium verticillioides include two required for microconidial chain formation. Fungal Genet. Biol. 41:852–864 Available at: https://www.sciencedirect.com/science/article/pii/S1087184504000672 [Accessed March 26, 2019].

15. Gale, L. R., Harrison, S. A., Ward, T. J., O’Donnell, K., Milus, E. A., Gale, S. W., et al. 2011. Nivalenol-type populations of Fusarium graminearum and F. asiaticum are prevalent on wheat in southern Louisiana. Phytopathology. 101:124–134 Available at: http://apsjournals.apsnet.org/doi/10.1094/PHYTO-03-10-0067 [Accessed March 26, 2018].

16. Gardiner, D. M., Osborne, S., Kazan, K., and Manners, J. M. 2009. Low pH regulates the production of deoxynivalenol by Fusarium graminearum. Microbiology. 155:3149–3156.

17. Gardiner, D. M., Stephens, A. E., Munn, A. L., and Manners, J. M. 2013. An ABC pleiotropic drug resistance transporter of Fusarium graminearum with a role in crown and root diseases of wheat. FEMS Microbiol. Lett. 348:36–45.

18. Garreau de Loubresse, N., Prokhorova, I., Holtkamp, W., Rodnina, M. V., Yusupova, G., and Yusupov, M. 2014. Structural basis for the inhibition of the eukaryotic ribosome. Nature. 513:517–522 Available at: http://dx.doi.org/10.1038/nature13737 [Accessed January 18, 2018].

19. Goswami, R. S. 2012. Targeted Gene Replacement in Fungi Using a Split-Marker Approach. Methods Mol. Biol. 835:255–269 Available at: http://link.springer.com/10.1007/978-1-61779-501-5 [Accessed May 8, 2019].

20. Goswami, R. S., and Kistler, H. C. 2005. Pathogenicity and in planta mycotoxin accumulation among members of the Fusarium graminearum species complex on wheat and rice. Phytopathology. 95:1397–1404.

21. Gulshan, K., and Moye-Rowley, W. S. 2007. Multidrug resistance in fungi. Eukaryot. Cell. 6:1933–1942.

22. Harris, L. J., and Gleddie, S. C. 2001. A modified Rpl3 gene from rice confers tolerance of the Fusarium graminearum mycotoxin deoxynivalenol to transgenic tobacco. Physiol. Mol. Plant Pathol. 58:173–181.

23. Hothorn, T., Bretz, F., and Westfall, P. 2008. Simultaneous inference in general parametric models. Biometrical J. 50:346–363.

24. Imazaki, I., and Kadota, I. 2015. Molecular phylogeny and diversity of Fusarium endophytes isolated from tomato stems. FEMS Microbiol. Ecol. 91:fiv098 Available at: http://www.ncbi.nlm.nih.gov/pubmed/26298015 [Accessed March 26, 2019].

25. Keller, N. P., Turner, G., and Bennett, J. W. 2005. Fungal secondary metabolism — from biochemistry to genomics. Nat. Rev. Microbiol. 3:937–947 Available at: http://www.nature.com/doifinder/10.1038/nrmicro1286.

26. Kettle, A. J., Batley, J., Benfield, A. H., Manners, J. M., Kazan, K., and Gardiner, D. M. 2015. Degradation of the benzoxazolinone class of phytoalexins is important for virulence of Fusarium pseudograminearum towards wheat. Mol. Plant Pathol. 16:946–962 Available at: http://doi.wiley.com/10.1111/mpp.12250 [Accessed June 26, 2018].

27. Klittich, C., and Leslie, J. F. 1988. Nitrate reduction mutants of Fusarium moniliforme (Gibberella fujikuroi). Genetics. 118:417–23 Available at: https://www.ncbi.nlm.nih.gov/pmc/articles/PMC1203296/pdf/ge1183417.pdf [Accessed October 22, 2019].

28. Koch, A., Biedenkopf, D., Furch, A., Weber, L., Rossbach, O., Abdellatef, E., et al. 2016. An RNAi-based control of Fusarium graminearum infections through spraying of long dsRNAs involves a plant passage and is controlled by the fungal silencing machinery ed. Savithramma P. Dinesh-Kumar. PLoS Pathog. 12:e1005901 Available at: http://dx.plos.org/10.1371/journal.ppat.1005901 [Accessed May 15, 2018].

29. Kovalchuk, A., and Driessen, A. J. 2010. Phylogenetic analysis of fungal ABC transporters. BMC Genomics. 11:177 Available at: http://bmcgenomics.biomedcentral.com/articles/10.1186/1471-2164-11-177 [Accessed November 20, 2018].

30. Lee, S., Son, H., Lee, J., Lee, Y. R., and Lee, Y. W. 2011. A putative ABC transporter gene, ZRA1, is required for zearalenone production in Gibberella zeae. Curr. Genet. 57:343–351.

31. Lofgren, L. A., LeBlanc, N. R., Certano, A. K., Nachtigall, J., LaBine, K. M., Riddle, J., et al. 2018. Fusarium graminearum: pathogen or endophyte of North American grasses? New Phytol. 217:1203–1212 Available at: http://doi.wiley.com/10.1111/nph.14894 [Accessed April 18, 2018].

32. Ma, Z., and Michailides, T. J. 2005. Advances in understanding molecular mechanisms of fungicide resistance and molecular detection of resistant genotypes in phytopathogenic fungi. Crop Prot. 24:853–863.

33. Menke, J., Dong, Y., and Kistler, H. C. 2012. Tri12p Influences virulence to wheat and trichothecene accumulation. Mol. Plant-Microbe Interact. 25:1408–1418.

34. Menke, J., Weber, J., Broz, K., and Kistler, H. C. 2013. Cellular development associated with induced mycotoxin synthesis in the filamentous fungus Fusarium graminearum. PLoS One. 8:e63077.

35. Nakajima, Y., Koseki, N., Sugiura, R., Tominaga, N., Maeda, K., Tokai, T., et al. 2015. Effect of disrupting the trichothecene efflux pump encoded by FgTri12 in the nivalenol chemotype of Fusarium graminearum. J. Gen. Appl. Microbiol. 61:93–96 Available at: https://www.jstage.jst.go.jp/article/jgam/61/3/61_93/_article.

36. O’Mara, S. P., Broz, K., Boenisch, M., Zhong, Z., Dong, Y., and Corby Kistler, H. 2020. The Fusarium graminearum t-SNARE Sso2 is involved in growth, defense, and DON accumulation and virulence. Mol. Plant-Microbe Interact. 33:888–901.

37. Oide, S., Berthiller, F., Wiesenberger, G., Adam, G., and Turgeon, B. G. 2014. Individual and combined roles of malonichrome, ferricrocin, and TAFC siderophores in Fusarium graminearum pathogenic and sexual development. Front. Microbiol. 5:759 Available at: http://journal.frontiersin.org/article/10.3389/fmicb.2014.00759/abstract [Accessed May 20, 2020].

38. Pasquali, M., and Kistler, C. 2006. Gibberella zeae ascospore production and collection for microarray experiments. J. Vis. Exp. 1:115 Available at: http://www.ncbi.nlm.nih.gov/pubmed/18704186 [Accessed March 26, 2018].

39. Perlin, M. H., Andrews, J., and Toh, S. S. 2014. Essential letters in the fungal alphabet: ABC and MFS transporters and their roles in survival and pathogenicity. Elsevier. Available at: http://dx.doi.org/10.1016/B978-0-12-800271-1.00004-4.

40. Pestka, J. J. 2007. Deoxynivalenol: Toxicity, mechanisms and animal health risks. Anim. Feed Sci. Technol. 137:283–298 Available at: http://linkinghub.elsevier.com/retrieve/pii/S0377840107002209.

41. Poppenberger, B., Berthiller, F., Lucyshyn, D., Sieberer, T., Schuhmacher, R., Krska, R., et al. 2003. Detoxification of the Fusarium mycotoxin deoxynivalenol by a UDP-glucosyltransferase from Arabidopsis thaliana. J. Biol. Chem. 278:47905–47914 Available at: http://www.jbc.org/lookup/doi/10.1074/jbc.M307552200.

42. Proctor, R. H. 1995. Reduced virulence of Gibberella zeae caused by disruption of a trichothecene toxin biosynthetic gene. Mol. Plant-Microbe Interact. 8:593 Available at: http://www.ncbi.nlm.nih.gov/pubmed/8589414 [Accessed October 2, 2019].

43. Proctor, R. H., Brown, D. W., Plattner, R. D., and Desjardins, A. E. 2003. Co-expression of 15 contiguous genes delineates a fumonisin biosynthetic gene cluster in Gibberella moniliformis. Fungal Genet. Biol. 38:237–249.

44. Proctor, R. H., McCormick, S. P., Alexander, N. J., and Desjardins, A. E. 2009. Evidence that a secondary metabolic biosynthetic gene cluster has grown by gene relocation during evolution of the filamentous fungus Fusarium. Mol. Microbiol. 74:1128–1142.

45. Puhalla, J. E., and Spieth, P. T. 1983. Heterokaryosis in Fusarium moniliforme. Exp. Mycol. 7:328–335 Available at: https://www.sciencedirect.com/science/article/pii/0147597583900178 [Accessed October 22, 2019].

46. Qi, P. F., Zhang, Y. Z., Liu, C. H., Zhu, J., Chen, Q., Guo, Z. R., et al. 2018. Fusarium graminearum ATP-binding cassette transporter gene FgABCC9 is required for its transportation of salicylic acid, fungicide resistance, mycelial growth and pathogenicity towards wheat. Int. J. Mol. Sci. 19.

47. Qi, T., Guo, Jia, Peng, H., Liu, P., Kang, Z., and Guo, Jun. 2019. Host-induced gene silencing: A powerful strategy to control diseases of wheat and barley. Int. J. Mol. Sci. 20:206 Available at: http://www.mdpi.com/1422-0067/20/1/206 [Accessed April 17, 2019].

48. R Core Team. 2018. R: A language and environment for statistical computing. Available at: https://www.r-project.org/.

49. Rocha, O., Ansari, K., and Doohan, F. M. 2005. Effects of trichothecene mycotoxins on eukaryotic cells: A review. Food Addit. Contam. 22:369–378 Available at: http://www.tandfonline.com/doi/abs/10.1080/02652030500058403.

50. Rodriguez Estrada, A. E., Hegeman, A., Corby Kistler, H., and May, G. 2011. In vitro interactions between Fusarium verticillioides and Ustilago maydis through real-time PCR and metabolic profiling. Fungal Genet. Biol.

51. Sanglard, D. 2016. Emerging threats in antifungal-resistant fungal pathogens. Front. Med. 3:1 Available at: http://www.ebi.ac.uk/interpro/entry/IPR011701/ [Accessed September 23, 2020].

52. Seong, K.-Y. Y., Pasquali, M., Zhou, X., Song, J., Hilburn, K., McCormick, S., et al. 2009. Global gene regulation by Fusarium transcription factors Tri6 and Tri10 reveals adaptations for toxin biosynthesis. Mol. Microbiol. 72:354–367 Available at: http://onlinelibrary.wiley.com/store/10.1111/j.1365-2958.2009.06649.x/asset/j.1365-2958.2009.06649.x.pdf?v=1&t=hv4dt99w&s=f7b520d8ee2c82b3abbb3cb2768a01898c6ba636.

53. Sieber, C. M. K., Lee, W., Wong, P., Münsterkötter, M., Mewes, H. W., Schmeitzl, C., et al. 2014. The Fusarium graminearum genome reveals more secondary metabolite gene clusters and hints of horizontal gene transfer. PLoS One. 9:e110311 Available at: http://www.ncbi.nlm.nih.gov/pubmed/25333987 [Accessed November 8, 2019].

54. Wachowska, U., Packa, D., and Wiwart, M. 2017. Microbial inhibition of Fusarium pathogens and biological modification of trichothecenes in cereal grains. Toxins (Basel). 9:408 Available at: http://www.ncbi.nlm.nih.gov/pubmed/29261142 [Accessed March 26, 2019].

55. Wang, Q., Chen, D., Wu, M., Zhu, J., Jiang, C., Xu, J.-R., et al. 2018. MFS Transporters and GABA Metabolism Are Involved in the Self-Defense Against DON in Fusarium graminearum. Front. Plant Sci. 9:438 Available at: http://journal.frontiersin.org/article/10.3389/fpls.2018.00438/full [Accessed April 13, 2018].

56. Waweru, B., Turoop, L., Kahangi, E., Coyne, D., and Dubois, T. 2014. Non-pathogenic Fusarium oxysporum endophytes provide field control of nematodes, improving yield of banana (Musa sp.). Biol. Control. 74:82–88 Available at: https://www.sciencedirect.com/science/article/pii/S1049964414000784 [Accessed March 26, 2019].

57. Wilson, W., Dahl, B., and Nganje, W. 2018. Economic costs of Fusarium Head Blight, scab and deoxynivalenol. World Mycotoxin J. 11:291–302 Available at: https://www.wageningenacademic.com/doi/10.3920/WMJ2017.2204 [Accessed January 14, 2019].

58. Wipfle, R., McCormick, S. P., Proctor, R. H., Teresi, J. M., Hao, G., Ward, T. J., et al. 2019. Synergistic phytotoxic effects of culmorin and trichothecene mycotoxins. Toxins (Basel). 11:555 Available at: https://www.mdpi.com/2072-6651/11/10/555 [Accessed May 12, 2020].

59. Yin, Y., Wang, Z., Cheng, D., Chen, X., Chen, Y., and Ma, Z. 2018. The ATP-binding protein FgArb1 is essential for penetration, infectious and normal growth of Fusarium graminearum. New Phytol. 219:1447–1466.

60. Zhang, X.-W., Jia, L.-J., Zhang, Y., Jiang, G., Li, X., Zhang, D., et al. 2012. In planta stage-specific fungal gene profiling elucidates the molecular strategies of Fusarium graminearum growing inside wheat coleoptiles. Plant Cell. 24:5159–76 Available at: http://www.pubmedcentral.nih.gov/articlerender.fcgi?artid=3556981&tool=pmcentrez&rendertype=abstract.

